# High-Throughput, Multiplex Immunofluorescence for the Computer-automated Immunophenotyping of Mycosis Fungoides

**DOI:** 10.64898/2025.12.30.697118

**Authors:** Patrick McMullan, Marc R. Benoit, Nathan Gasek, Sydney Riddick, Joseph Masison, Katalin Ferenczi, David Rowe, Gillian Weston

## Abstract

The diagnosis of Mycosis Fungoides (MF) is difficult and often delayed, exacerbated by the constraint of conventional immunohistochemistry (IHC) to analyze only one antigen per tissue section, often necessitating repeat biopsies and extensive workups. We sought to validate a high-throughput Multiplex Immunofluorescence (MIF) method, coupled with computer-automated image analysis, to generate comprehensive immunophenotyping data from a single formalin-fixed, paraffin-embedded (FFPE) biopsy. We applied an 11-biomarker MIF panel across 18 archived skin specimens (9 MF/TCR clonality positive and 9 control/TCR clonality negative). Initial validation confirmed that MIF antigen expression and spatial localization were concordant with sequential IHC-stained sections. Whole slide image stacks were analyzed using both computer-assisted and fully computer-automated pipelines. Both methods successfully delineated immunophenotypic differences. MF specimens showed a significant expansion of hematopoietic cells and proliferative T-lymphocytes compared to controls. Crucially, MF tissues also exhibited a significant increase in the percentage of atypical T-lymphocytes. Our results validate the potential of MIF to obtain comprehensive, high-dimensional diagnostic information from a single tissue section. Integration with computer-automated analysis offers a scalable, high-throughput platform that can significantly aid in the timely and accurate diagnosis of cutaneous lymphomas.

## Introduction

Mycosis Fungoides (MF) are a spectrum of non-Hodgkin lymphomas characterized by the expansion and accumulation of monoclonal malignant T-cells.^1,2^ Diagnosis of MF can be particularly challenging within the early stages of disease due to its equivocal clinical, and sometimes histopathologic presentation with an estimated mean delay in diagnosis being 2-4 years, and can require an estimated mean of 2.89 biopsies per patient.^3,4^ Histopathologic diagnosis of MF often relies on immunohistochemistry (IHC) staining to identify the surface antigen expression profile or “immunophenotype” of malignant monoclonal T-cells.^1,5^ While IHC is a primary and widely adopted modality for MF diagnosis, it is typically limited in its ability to visualize usually one, but rarely two antigens per single tissue section. This limitation can result in practical challenges especially within equivocal cutaneous lesions that can not only lead to exhaustion of the available tissue and need for repeat biopsies, but also often requires further diagnostic workup in the form of T-cell receptor (TCR) molecular studies and flow cytometry.^6–8^ In addition, these methods limit the potential to simultaneously visualize the comprehensive immunophenotype within the tumor microenvironment and readily depict between malignant or benign T-cells *in situ*. In an effort to obviate these shortcomings of conventional immunophenotyping with IHC, multiplex immunofluorescence (MIF) has emerged as a promising technology for the identification and visualization of multiple cellular populations within a single tissue section.^9–11^ MIF modalities can broadly be divided into simultaneous staining methodologies (i.e. antigens of interest are all labeled at once) as well as cyclical staining protocols.^12^ While simultaneous staining approaches are benefited by their relative simplicity, their throughput is limited by spectral overlap between staining fluorophores. To overcome this, cyclical methods utilize repeated cycles of staining and imaging tissue specimens with a panel of primary fluorescently-labeled antibodies that can be visualized by fluorescence microscopy (Supplemental Figure 1) followed by signal quenching.^9–11^ After each image acquisition, a photo-inactivation step can be performed to the stained tissue section to clear any remaining fluorescence signal from previous cycles that enables multiple subsequent staining cycles on the same section allowing for detection of >30 antigens.^9,13^ Concatenated digital whole slide image stacks of each specimen are generated to visualize the expression and spatial distribution of multiple antibodies and can be further processed computationally to generate comprehensive immunophenotyping data in similar quality to that of flow cytometry. Additionally, with generation of new computer-automated immunophenotyping pipelines, these microscopy data can be analyzed at the single cell resolution to distinguish the presence and relative abundance of distinct cellular populations between individual patient specimens in malignant versus benign disease states. Datasets can also be leveraged to explore spatial relationships among diverse cell populations that are unobtainable through traditional IHC or flow cytometry.

Multiplex immunofluorescence is an emerging dermatopathologic technique, particularly in the field of cutaneous oncology.^12,14–16^ Reports examining its utility in malignant melanoma and merkel cell carcinoma (MCC) demonstrated its ability to precisely characterize T-cell populations (CD3, CD8, FOXP3) and myeloid cells (CD68, CD163) alongside tumor markers (such as SOX2 for MCC or SOX10 for melanoma) and immune checkpoint molecules (PD-L1). This spatial analysis is crucial for predicting the response to checkpoint inhibitor immunotherapy in MCC and melanoma given that the proximity of cells like PD-1+ and PD-L1+ or cytotoxic CD8+ T-cells to tumor cells can be highly predictive of efficacy, a finding that surpasses single PD-L1 expression.^12,14–16^ MIF is also employed in hereditary epidermolysis bullosa within the context of immunofluorescence antigen mapping (IFM), recognized as a standard diagnostic modality that functions as a multiplex assay utilizing multiple structural protein antibodies on a single slide.^17,18^ The primary goal of this study was to examine the utility of an antibody-based MIF assay to identify, distinguish and quantify aberrant immune cell populations within MF skin tissue sections in comparison to control benign inflammatory dermatoses.^1,10,19,20^ Herein we first validated the relative expression and spatial localization of MIF stained tissue sections in comparison to sequential sections stained with conventional IHC and H&E. We subsequently demonstrate the potential to generate immunophenotyping data from MIF digital whole slide image stacks using two distinct image processing pipelines that collectively demonstrated significant variations in atypical T-cell populations between MF and control specimens.

## Results

### Validation of Multiplex Immunofluorescence Workflow

We first examined five IHC and TCR gene rearrangement confirmed MF cases to validate MIF as an effective immunophenotyping modality (Figure 1) by comparing relative tissue-antigen expression and spatial localization of MIF-stained sections to sequential H&E and conventional IHC-stained sections utilized to generate the initial MF diagnosis (Figure 1A-F). MIF provided concordant antigen expression and spatial localization for a multitude of antigens including CD4, CD5, CD7 and CD8 when compared to IHC (Figure 1G-L). Furthermore, in 80% (4/5) of examined cases, quantification of CD4:CD8 ratio from MIF whole slide image stacks directly correlated with the original ratio reported and subsequently quantified from conventional IHC (Figure 1M). MIF overestimated CD4:CD8 ratio as reported on the initial pathology report in 20% (1/5) of cases. Given this initial validation, an 11-biomarker panel frequently utilized in the diagnostic workup of MF (Table 2) was adapted to allow for four cycles of MIF on a single patient-derived tissue specimen (Figure 2A). This panel was used to examine 18 patient specimens with clinical and histopathologic suspicion for MF, consisting of 9 TCR clonality negative (“control”) cases as well as 9 TCR clonality positive (“MF”) cases. Processed whole slide digital image stacks demonstrated atypical monoclonal T-lymphocytes within the MF intraepidermal tumor microenvironment, with their immunophenotype exhibiting strong CD4 and CD45, marginal CD3 and CD5 and effective absence of CD7, CD8 and CD30 antigen expression (Figure 2B).^1^ Collectively, these data validate MIF and represent its potential to generate a highly comprehensive immunophenotyping dataset.

**Figure 1:**
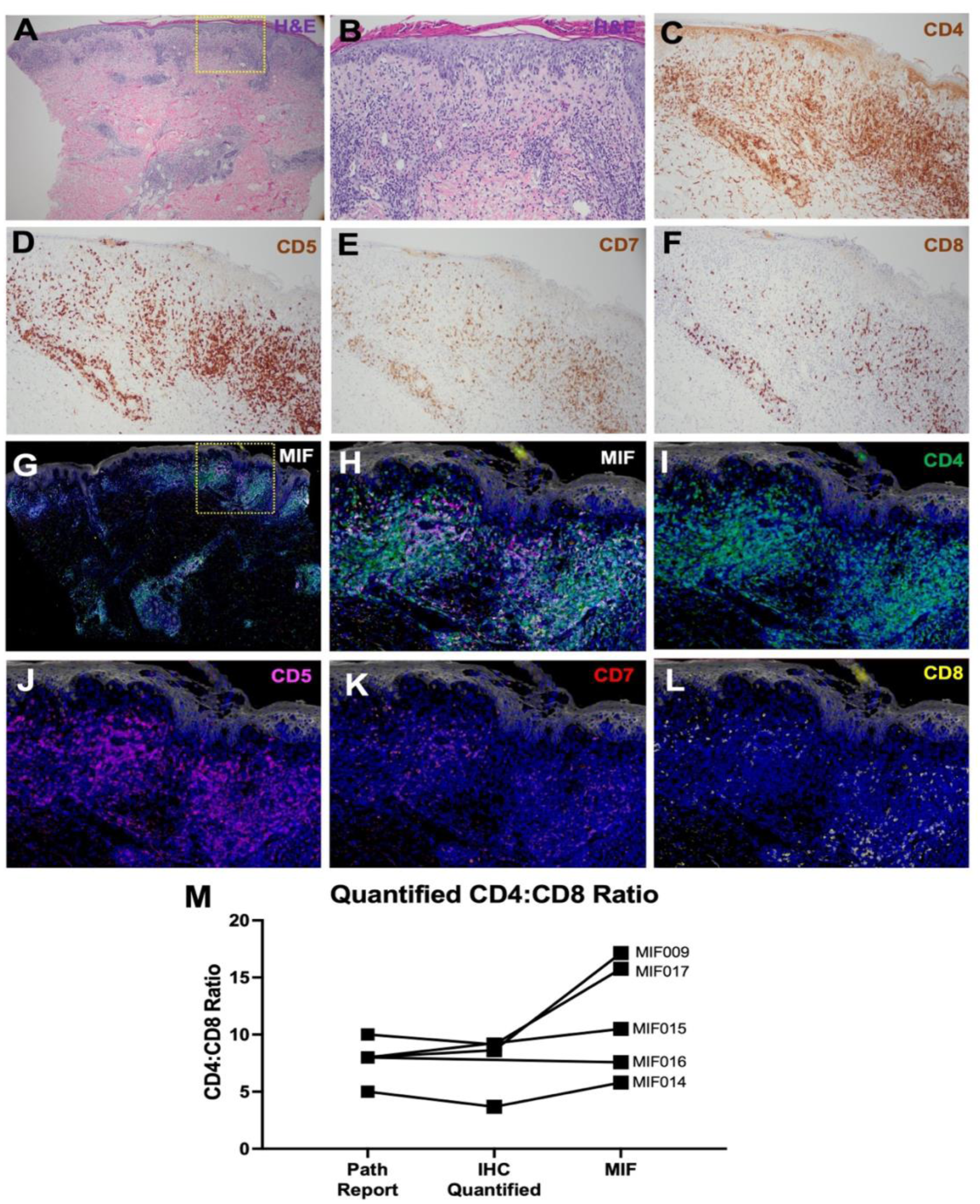
Representative images of (A-B) hematoxylin and eosin; (C-F) immunohistochemistry; and (G-L) multiplex immunofluorescence obtained from a single MF tissue specimen. Note the similarities in both degree of antigen expression and spatial localization of target antigens between serial tissue sections stained using conventional immunohistochemistry versus a single tissue section following multiplex immunofluorescence imaging. (M) Line graph comparing the quantified CD4:CD8 ratio among MF cases on the basis of reported ratio by the reading dermatopathologist (Path Report) or computer-assisted quantification of conventional IHC stained specimens (IHC quantified) or multiplex immunofluorescence stained specimens (MIF).

**Figure 2:**
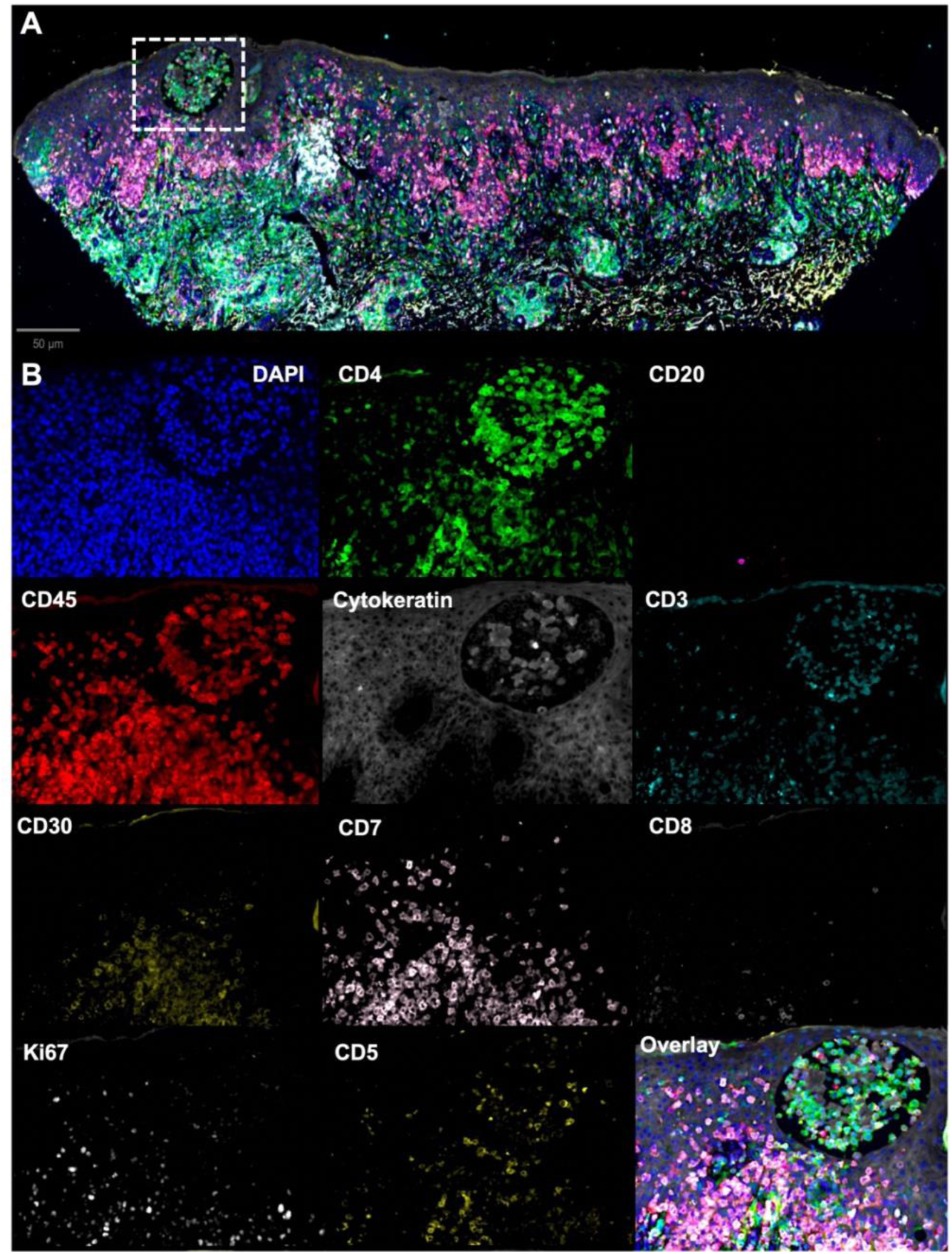
Representative (A) low- and (B) high magnification of a concatenated digital image stack generated from a MF patient specimen following 4 cycles of antibody-based multiplex immunofluorescence. Individual antibodies are labeled within each high magnification figure, and a concatenated overlay image is present

### Comprehensive Immunophenotyping by MIF reveals global distinctions in cellular populations between MF and control samples

MIF whole slide image stacks were subsequently quantified using the open-source image cytometry software QuPath. Following cell detection and segmentation, automated thresholding was used for each individual antibody within the MIF panel^21^ to discern variations in global immunophenotyping between control and MF specimens (Figures 3A-D). MF specimens exhibited a significant reduction in the relative abundance of pan-cytokeratin positive epithelial cells and displayed a significant expansion in CD45+ hematopoietic cell populations, CD4+ T-lymphocytes and Ki67+ proliferative cells when compared to control specimens (Figure 3C,D). In addition to demonstrating global immunophenotypic variations, we quantified critical parameters relied upon for MF diagnostic workup including: (1) CD4:CD8 T-lymphocyte ratio; (2) atypical malignant CD4 T-lymphocytes percentage according to surface expression profile of CD45+,CD4+,CD5- or CD45+,CD4+,CD7- expression; and (3) proliferative index demonstrating the percentage of Ki67+ cell types (Figure 3E-G). While we did not identify significant variations within the calculated CD4:CD8 ratio, MF specimens globally displayed a significant increase in the number of atypical CD45+,CD4+,CD7- T-lymphocytes and proliferative CD4+ T-lymphocytes (Figure 3E-G). Secondary analyses performed on intraepidermal immune populations did not reveal significant variations in CD4:CD8 ratio or atypical T-lymphocyte percentage; however, MF specimens similarly revealed an increase in the percentage of intraepithelial Ki67+ cells (Supplemental Figure 2). Collectively, these data provide direct evidence demonstrating the potential of MIF image stacks to delineate distinct immunophenotypic profiles among clinically and histopathologically equivocal MF and control specimens.

**Figure 3:**
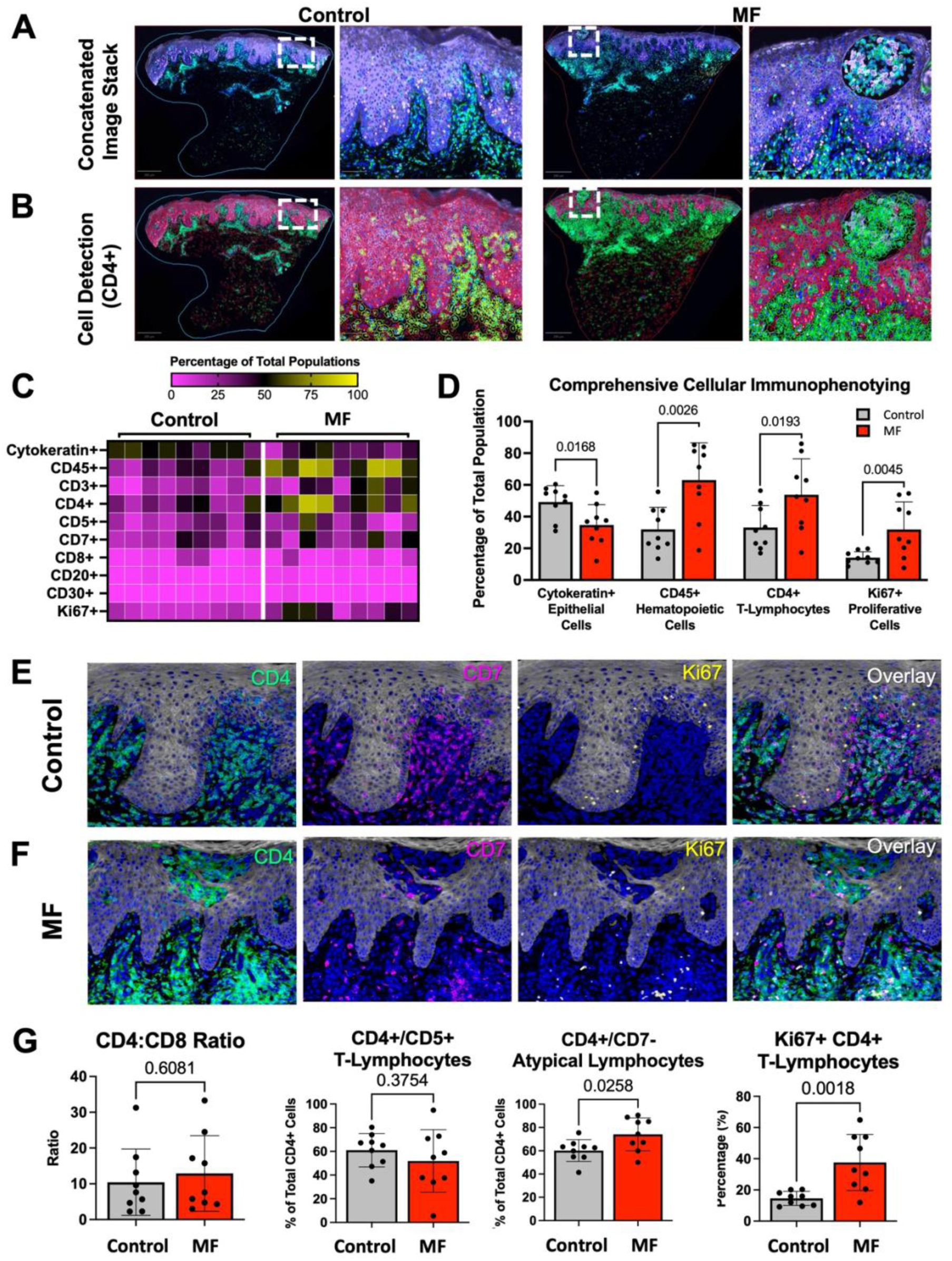
(A-B) Image stacks from (A) control and (B) MF patient specimens (top) demonstrating expression of CD4 (green), Ki67 (orange) and pan-cytokeratin (purple). Cell detection and image analysis workflow employed on control and MF concatenated image stacks (bottom) depict cell expression thresholds for CD4+ populations (green) versus CD4- populations (pink). (C) Heatmap of comprehensive immunophenotyping data from control and MF specimens. (D) Bar graph of comprehensive immunophenotyping data obtained from control (n=9) and MF (n=9) patient image stacks. (E-F) Representative high power MIF images of (E) control and (F) MF patient specimens demonstrating expression of CD4 (green), CD7 (pink), Ki67 (yellow) with a concatenated overlay. (G) Bar graphs of quantified CD4:CD8 ratio, percentage of CD45+, CD4+,CD5+, percentage of atypical lymphocytes (CD45+,CD4+,CD7-) and proliferative (Ki67+) CD4+ T-lymphocytes within control (n=9) and MF (n=9) patient image stacks.

### Processing of MIF stacks using a computer-automated image cytology pipeline readily identified immunophenotype distinctions between control and MF specimens

Encouraged by our initial immunophenotying data generated using QuPath, we further explore the utility of an entirely computer-automated multiplex image analysis pipeline referred to as the steinbock toolkit^22^ to identify and quantify immunophenotyping data from MIF image stacks. Using both control and MF image stacks as the primary input, the steinbock toolkit effectively immunophenotyped 12 distinct cellular populations according to antibody expression by using its cell segmentation and hierarchical clustering algorithms (Figure 4A,B and Supplemental Figure 3). Further analyses of these populations within MF samples revealed a significant reduction in the total number of pan-cytokeratin positive keratinocytes, and a significant increase in the number of atypical CD4+, CD7- T-lymphocytes when compared to controls (Figure 4C). These computer-automated data not only directly correlate with the previous computer-assisted data, but they also demonstrate, for the first time to our knowledge, the successful use of a computer-automated MIF image analysis pipeline to aid in the diagnosis of cutaneous lymphoma in a potentially high-throughput manner.

**Figure 4:**
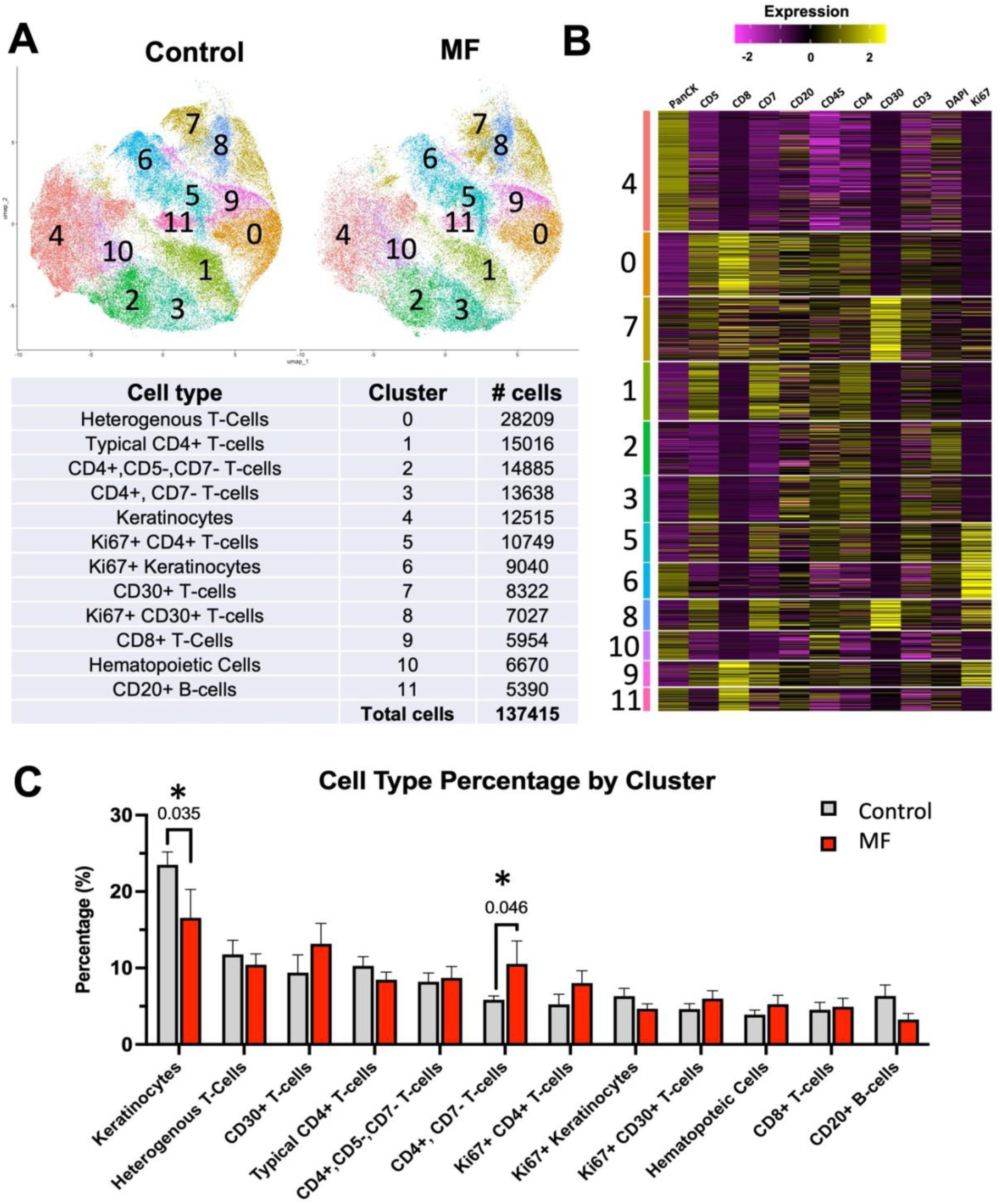
(A) Representative UMAP plot demonstrating the 12 distinct cellular populations identified using the steinbock toolkit and their associated cell types. (B) Representative heatmap depicting varying surface marker expression among cell types. (C) Bar graphs of comprehensive immunophenotyping data obtained from control and MF image stacks demonstrating relative percentage of each cell type across the entire number of analyzed cells. Clusters with significant variations between control and MF specimens identified with *p* value.

## Discussion

Cutaneous T-Cell Lymphomas encompass a rare spectrum of skin cancers that can be difficult to diagnose due to the need for extensive cellular and molecular diagnostic tests. Here we demonstrate the potential of MIF to obtain high-dimensionality immunophenotyping information from a single skin tissue section in a condition that often requires multiple skin tissue sections and/or biopsies. We specifically examined equivocal cutaneous lesions with both clinical and histologic features concerning for MF that required additional diagnostic workup in the form of both IHC and TCR gene rearrangement. Herein, we developed an 11-antibody panel that could distinguish distinct immunophenotypic signatures between control and MF specimens. We successfully identified a population of atypical lymphocytes within MF specimens defined by a CD45+,CD4+,CD7- immunophenotype that are significantly enriched compared to control tissues.^23^ Given that immunophenotyping via MIF based assays may help predict response to therapies such as immune checkpoint inhibitors,^24^ this work may serve as a foundation for future efforts in precision medicine.

While these data support the use of MIF in the diagnosis of cutaneous lymphomas, we greatly appreciate the limitations of this current antibody panel given the breadth of lymphoproliferative disorders and associated antigenic variation. This can be addressed in future studies by implementing additional antigens within or panel including but not limited to CD56, βF1, TCR𝛾/𝛿, PD1, ALK, or EMA ^25^ to characterize aberrant B- and T-cell neoplasms. We also acknowledge there are several technical limitations to our MIF protocol in its current form that restrict widespread adoption into clinical pathology and laboratory medicine. Perhaps one of the greatest technical hurdles includes access to capital equipment including a multi-channel fluorescent microscope and camera for image acquisition. Additionally, workflow standardization with respect to tissue specimen antigen retrieval, primary antibody incubation, and photoinactivation treatment is a significant technical hurdle to more universal adoption. Considering these hurdles, there is a growing effort to not only validate antibodies from research applications suitable for MIF but also detail standardized protocols and reagents that could be implemented into an automated slide staining device workflow.^9,13,26^ Within our described protocol, one technical hurdle in utilizing an automated slide staining device is the use of a photoinactivation step that may preclude a completely automated workflow in a traditional pathology laboratory. One final limitation includes proficiency in image processing and analysis of digital whole slide image stacks. Here, we have utilized two validated image analysis pipelines including QuPath for computer-assisted analysis in addition to steinbock which is a fully automated image analysis workflow that requires technical expertise in bioinformatics and statistical software. There are ongoing efforts to utilize artificial intelligence and develop computer-automated image analysis software to efficiently process and analyze MIF images.^15,27,28^

In conclusion, our data can serve as an essential framework demonstrating the potential of multiplex imaging in improving the diagnosis of skin cancers and inflammatory skin conditions. This will provide a foundation for the development of additional disease-specific MIF panels and imaging technologies.

## Materials and Methods

### Human Biospecimen Selection

All studies were approved by the University of Connecticut Health Center’s Institutional Review Board (Protocol #23-137-1). Archived formalin-fixed, paraffin embedded (FFPE) biospecimens were identified by retrospective review of pathology reports that included the term “mycosis fungoides.” Biospecimens were selected that included the following criteria: (1) cases with a differential diagnosis of mycosis fungoides; AND (2) undergone IHC for diagnostic workup; AND (3) sent for molecular studies in the form of TCR gene rearrangement. Eighteen biospecimens were selected and sectioned for downstream MIF analyses (Table 1) that included 9 spongiotic dermatitis cases with negative TCR gene clonality categorized as “control” biospecimens and 9 TCR gene clonality positive cases categorized as “MF” biospecimens.

**Table 1:** Selected Biospecimen Characteristics.

| **Study Identifier** | **TCR Gene Clonality** | **Rendered Histopathologic Diagnosis** | **Peripheral Blood CD4:CD8 Ratio (If Obtained)** |
| --- | --- | --- | --- |
| MIF000 | Negative | Spongiotic Dermatitis | 3.9 : 1.0 |
| MIF001 | Negative | Spongiotic Dermatitis | 3.9 : 1.0 |
| MIF002 | Negative | Psoriasiform spongiotic dermatitis with eosinophils | 3.9 : 1.0 |
| MIF003 | Negative | Psoriasiform dermatitis with eosinophils | - |
| MIF004 | Negative | Psoriasiform spongiotic dermatitis with focal lymphocyte exocytosis | - |
| MIF005 | Negative | Spongiotic dermatitis | - |
| MIF006 | Negative | Psoriasiform spongiotic dermatitis | - |
| MIF007 | Negative | Psoriasiform dermatitis with eosinophils | - |
| MIF008 | Negative | Spongiotic dermatitis with focal interface dermatitis | - |
| MIF009 | Positive | Mycosis fungoides, Lymphocyte exocytosis, focal spongiosis and parakeratosis | 1.6 : 1.0 |
| MIF010 | Positive | Mycosis fungoides, Lymphocyte exocytosis with lymphocyte atypia and superficial perivascular lymphocyte infiltrate | 1.8 : 1.0 |
| MIF011 | Positive | Mycosis fungoides, Lichenoid dermatitis with prominent lymphocyte exocytosis | 2.8 : 1.0 |
| MIF012 | Positive | Mycosis fungoides, Atypical lymphocytic exocytosis | - |
| MIF013 | Positive | Mycosis fungoides, Interface dermatitis with lymphocyte exocytosis | 1.6 : 1.0 |
| MIF014 | Positive | Mycosis fungoides, Atypical epidermotropic lymphocytic infiltrate | - |
| MIF015 | Positive | Mycosis fungoides, Prominent lymphocyte exocytosis and superficial and mid-dermal perivascular lymphocytic infiltrate | 3.8 : 1.0 |
| MIF016 | Positive | Mycosis fungoides, Lymphocyte exocytosis with lymphocyte atypia and superficial perivascular lymphocytic infiltrate | - |
| MIF017 | Positive | Mycosis fungoides, Lymphocyte exocytosis with formation of intraepidermal microabscesses | - |

### Slide Preparation and Antigen Retrieval

5µm FFPE sections were first deparaffinized and rehydrated with xylene and graded alcohol treatments and subsequently underwent heat-based sodium citrate (pH 6.0) antigen retrieval for 15 minutes at 95*C. Sections were washed three times with phosphate buffered saline and pre-treated with photobleaching solution (containing 4.5% hydrogen peroxide and 100 mM sodium hydroxide) for 1 hour while exposed to white light to reduce background autofluorescence. ^9,26,29^

### Multiplex Immunofluorescence

Slides were permeabilized in 0.1% Triton X for 10 minutes, blocked using 2% bovine serum albumin in PBS for 45 minutes and stained with fluorescently-conjugated primary antibody for 1 hour at room temperature (antibodies and dilutions listed in Table 2). The slides were subsequently washed in PBS and mounted in 50% glycerol/PBS solution containing 4’,6-diamidino-2-phenylindole (DAPI) (1:1000 dilution) and imaged using a Zeiss Axioscan Z1 high-speed automated image acquisition system (Cat#440640-9903-000) and a high resolution camera (AxioCam HRm). Slide visualization was completed using filter cubes for 4 standard fluorescent channels (DAPI/Alexa350, FITC/Alexa488, Texas Red/Alexa594 and CY5/Alexa647). Following image acquisition, slides were submerged in PBS to allow for coverslip removal without compromising tissue integrity and treated with photobleaching solution again for 60 minutes while exposed to white light. This cyclic process continued for a total of 4 rounds of MIF imaging.

**Table 2:** Antibodies Utilized for Multiplex Immunofluorescence.

| **Table 2: Antibodies Utilized for Multiplex Immunofluorescence** |  |  |
| --- | --- | --- |
| **<u>Antibody</u>** | **<u>Source</u>** | **<u>Dilution Factor</u>** |
| **CD3 Alexa Fluor® 546-conjugated Antibody**<br>**Mouse Monoclonal** | Santa Cruz<br>(PC3/188A) | 1:100 |
| **CD4 Alexa Fluor® 488-conjugated Antibody**<br>**Polyclonal Goat IgG** | R&D Systems FAB8165G | 1:100 |
| **CD5 CF® 594-conjugated Antibody**<br>**Mouse Monoclonal** | Biotium (CD5/2416) | 1:100 |
| **CD7 CoraLite® 594-conjugated Antibody**<br>**Mouse Monoclonal** | ThermoFisher (1G10D8) | 1:200 |
| **CD8 CF® 647-conjugated Antibody**<br>**Mouse Monoclonal** | Biotium (C8/468) | 1:100 |
| **CD20 CF® 647-conjugated Antibody**<br>**Mouse Monoclonal** | Biotium (C20/MS4A1) | 1:100 |
| **CD30 CF® 647-conjugated Antibody**<br>**Mouse Monoclonal** | Biotium<br>(CD30 (Ki-1/779)) | 1:100 |
| **CD45 CF® 488A-conjugated Antibody**<br>**Mouse Monoclonal** | Biotium (CD45 /<br>LCA(2B11+PD7/26)) | 1:100 |
| **CD47 CF® 488A-conjugated Antibody**<br>**Mouse Monoclonal** | Biotium (CD47/2937) | 1:100 |
| **Ki67 CF® 488A-conjugated Antibody**<br>**Mouse Monoclonal** | Biotium (MKI67/2462) | 1:100 |
| **Pan-Cytokeratin CF® 647-conjugated Antibody**<br>**Mouse Monoclonal** | Biotium<br>(KRTL/1077+KRTH/1076) | 1:300 |

### Image Stack Assembly

Digital whole slide image stacks were uploaded, concatenated and aligned using the nuclear (DAPI) channel as a common reference marker between subsequent imaging rounds by using the HyperStackReg plugin^30^ within the open source software ImageJ.

### Cell detection, segmentation and quantification of single cell fluorescence intensity

Concatenated digital image stacks were first uploaded to the open source bioimage analysis software QuPath.^21^ For images obtained by fluorescence microscopy, cell segmentation and detection was performed using the nuclear (DAPI) channel for fluorescence channels and hematoxylin for brightfield images. Quantification of cell number for each individual channel (marker) was identified and employed across all samples using the automated cell detection feature. Quantification of cells expressing multiple markers was completed using the composite classifier function. Comprehensive immunophenotyping of whole slide images was determined by examining the total number of cells expressing individual surface markers in comparison to the total number of cells within the entire image. Phenotyping of intraepidermal immune populations was generated using a pixel classification modifier by identifying the total number of cells contained within a boarder of cytokeratin positive cells.

### Computer-automated multiplex image processing and analysis

For computer automated detection and immunophenotyping, a comprehensive end-to-end multiplex image analysis workflow referred to as the steinbock toolkit (developed by Windhager et al.)^22^ was implemented. The primary input consisted of digital whole slide concatenated image stacks under one image segmentation and pixel classification using a deep learning model incorporated into the workflow. The segmented cells and pixel classification data were further analyzed using both single cell and spatial analysis toolkits to allow for hierarchical clustering and heat map generation. For computational derived cellular clusters, manual inspection of immunophenotype data on basis of fluorescence intensity was completed to classify underlying cell etiology.

### Statistical analysis

All statistical analyses were performed using Graphpad Prism Version 9 (GraphPad Software, Inc., La Jolla, CA, USA) with *P*-values < 0.05 considered statistically significant. For all analyses that compared data obtained from control and MF cohorts in Figure 3 and Supplemental Figure 2 an unpaired two-tailed t-test was utilized. Additionally, all analyses comparing cell type percentages from computer-automated image processing in Figure 4 utilized a Wilcoxon rank sum test. Each *n* value refers to the number of biospecimens. All data points were included in the data analysis. No samples were excluded. *P*-values for each statistical comparison are indicated within the figure panels. All human data are included within the main and supplemental figures within this manuscript.

## Author Contributions

Patrick McMullan (Conceptualization, Data curation, Formal analysis, Funding Acquisition, Investigation, Methodology, Project Administration, Validation, Visualization, Writing—original draft preparation, Writing—review & editing), Marc R. Benoit (Methodology, Data curation, Formal analysis, Investigation, Validation, Visualization, Writing—review & editing), Nathan Gasek (Methodology, Data curation, Formal analysis, Investigation, Validation, Visualization, Writing—review & editing), Sydney Riddick (Methodology, Data curation, Formal analysis, Investigation, Validation, Visualization, Writing—review & editing), Joseph Masison (Data curation, Formal Analysis, Investigation, Validation, Writing—review & editing), Katalin Ferenczi (Project Administration, Methodology, Supervision, Validation, Writing—review) David W. Rowe (Funding Acquisition, Investigation, Project Administration, Methodology, Resources, Supervision, Validation, Writing—review & editing), and Gillian Weston (Conceptualization, Funding acquisition, Investigation, Project administration, Resources, Supervision, Validation, Visualization, Writing—original draft preparation, Writing—review & editing).

## IRB approval status

Approved

---

**Supplemental Figure 1:**
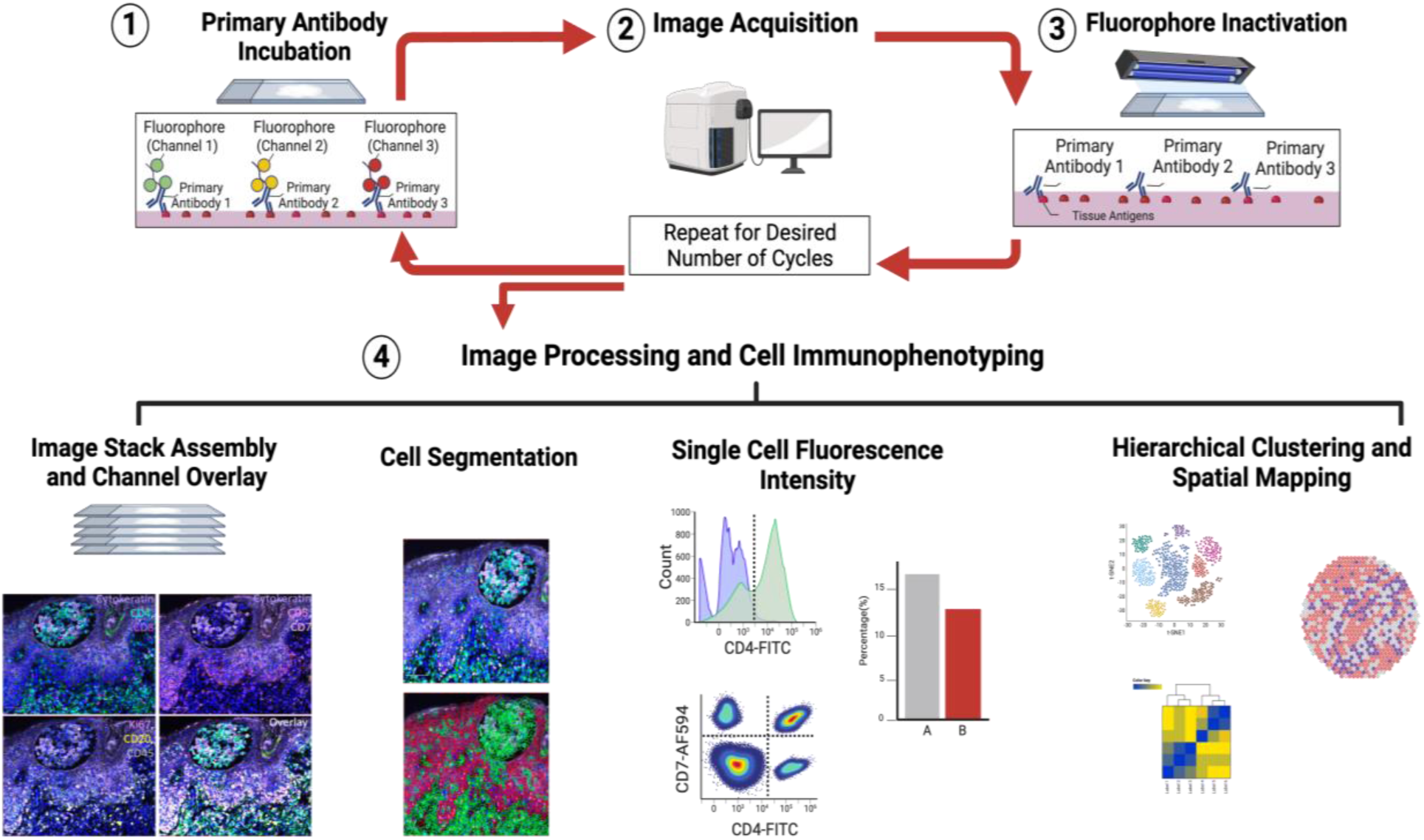
Representative flowchart of the multiplex immunofluorescence pipeline including both image acquisition, image stack assembly and the multiple formats of both computer-assisted and computer-automated image analysis.

**Supplemental Figure 2:**
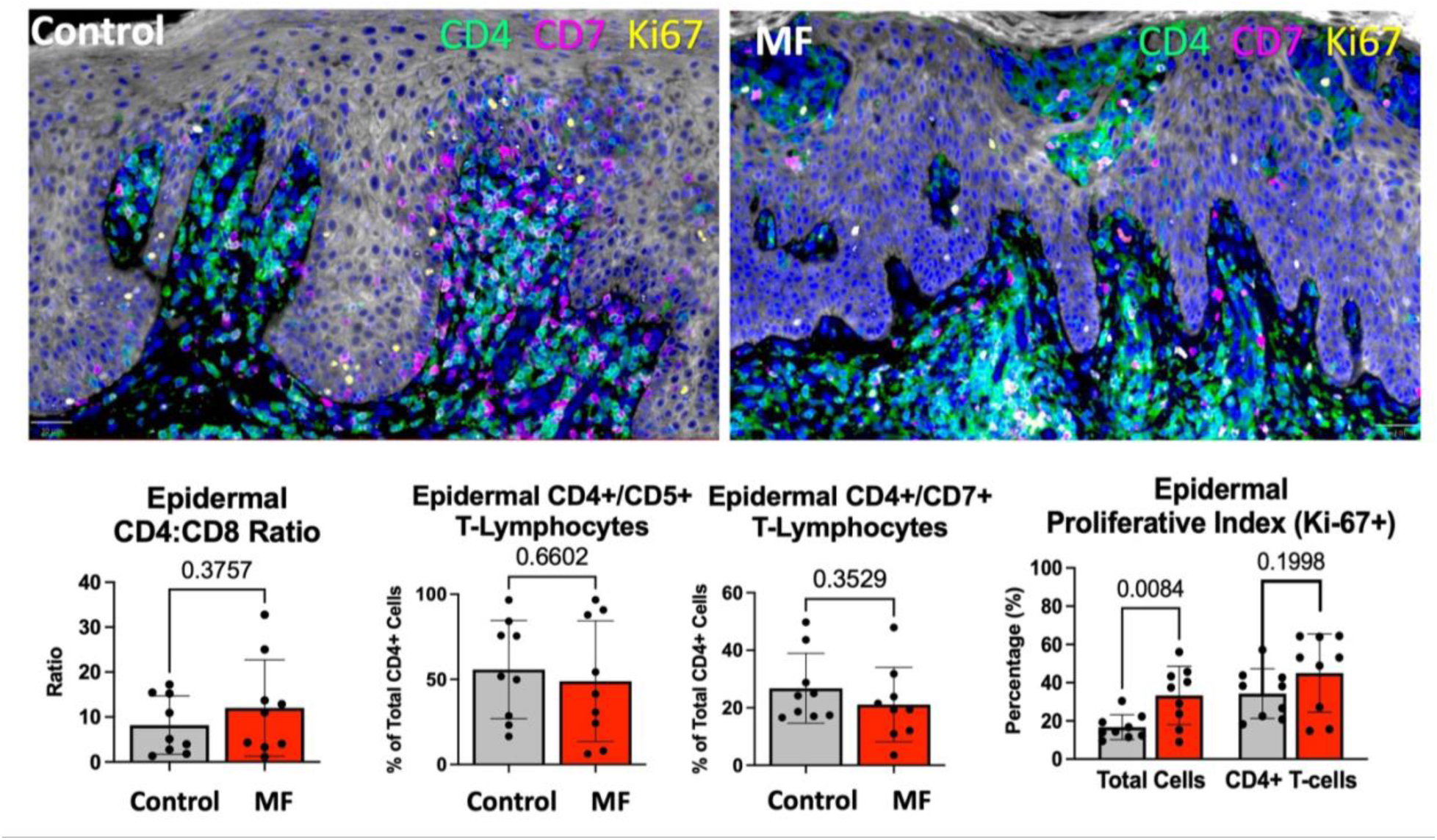
Representative concatenated high power MIF images of control and MF patient specimens demonstrating expression of CD4 (green), CD7 (pink), Ki67 (yellow). Bottom depicts Bar graphs of quantified CD4:CD8 ratio, percentage of CD45+, CD4+,CD5+, percentage of atypical lymphocytes (CD45+,CD4+,CD7-) and proliferative (Ki67+) CD4+ T-lymphocytes within control (n=9) and MF (n=9) patient image stacks.

**Supplemental Figure 3:**
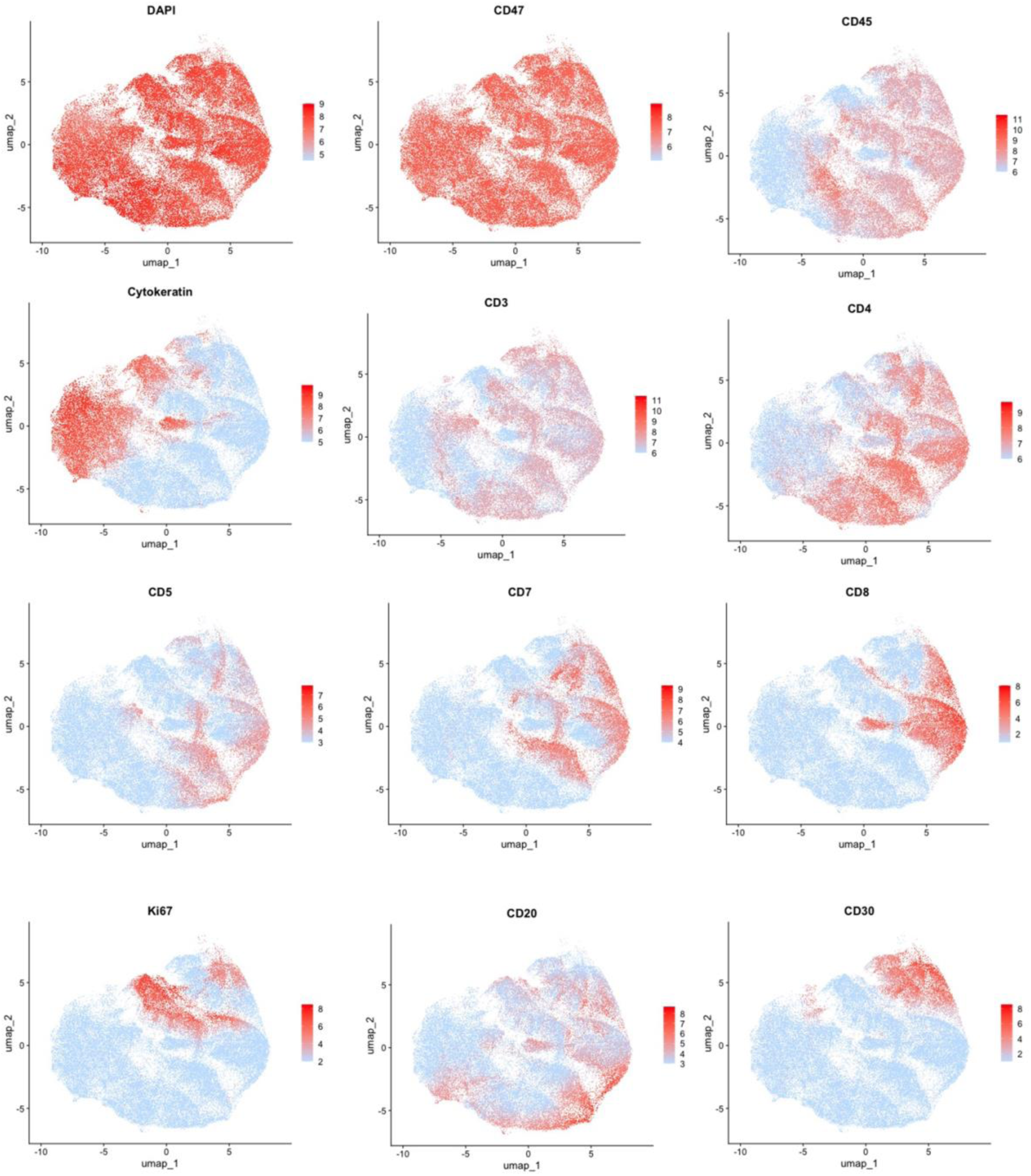
UMAP of all single segmented cells from combined control and MF MIF image stacks depicting relative spatial localization of antigen expression.

